# Rational redesign of antigen binding domain improves *in vivo* efficacy of CD22-CAR T cells

**DOI:** 10.1101/2025.03.13.643183

**Authors:** Kole R. DeGolier, Zachary H. Walsh, Lillie Leach, Christine Brzezinski, Amanda Novak, Dimiter Dmitrov, M. Eric Kohler, Terry J. Fry

## Abstract

Chimeric antigen receptor (CAR) T cells targeted to CD19 are an effective therapy for B-lineage malignancies. However, about half of patients relapse and this therapeutic, often with antigen-negative disease, warranting the targeting of other antigens. CD22 represents another promising target, with highly restricted but ubiquitous expression across the B-lineage. However, despite promising preclinical work by several groups with CD22-targeted CAR T cells targeting of this antigen in the clinic has proven difficult, with many patients relapsing with CD22^Lo^ leukemia, contrasting to complete loss of CD19 expression post CD19-CAR. While prior work has demonstrated that a CAR with so-called “tonic” antigen-independent signaling properties has proven to be highly efficacious, tonic signaling has been shown be detrimental to long-term T cell function. Here, we demonstrate a balance between binding affinity and antigen-independent tonic signaling (as determined by length of flexible linker) in determining CAR function. We show that maximal CAR function in the settings CD22^Lo^ and WT leukemia is maintained by boosting binding affinity without shortening flexible linker to induce tonic signaling, establishing rational modification of antigen binding domain as an important approach for modulating the function of cellular therapeutics.

**GRAPHICAL ABSTRACT:** 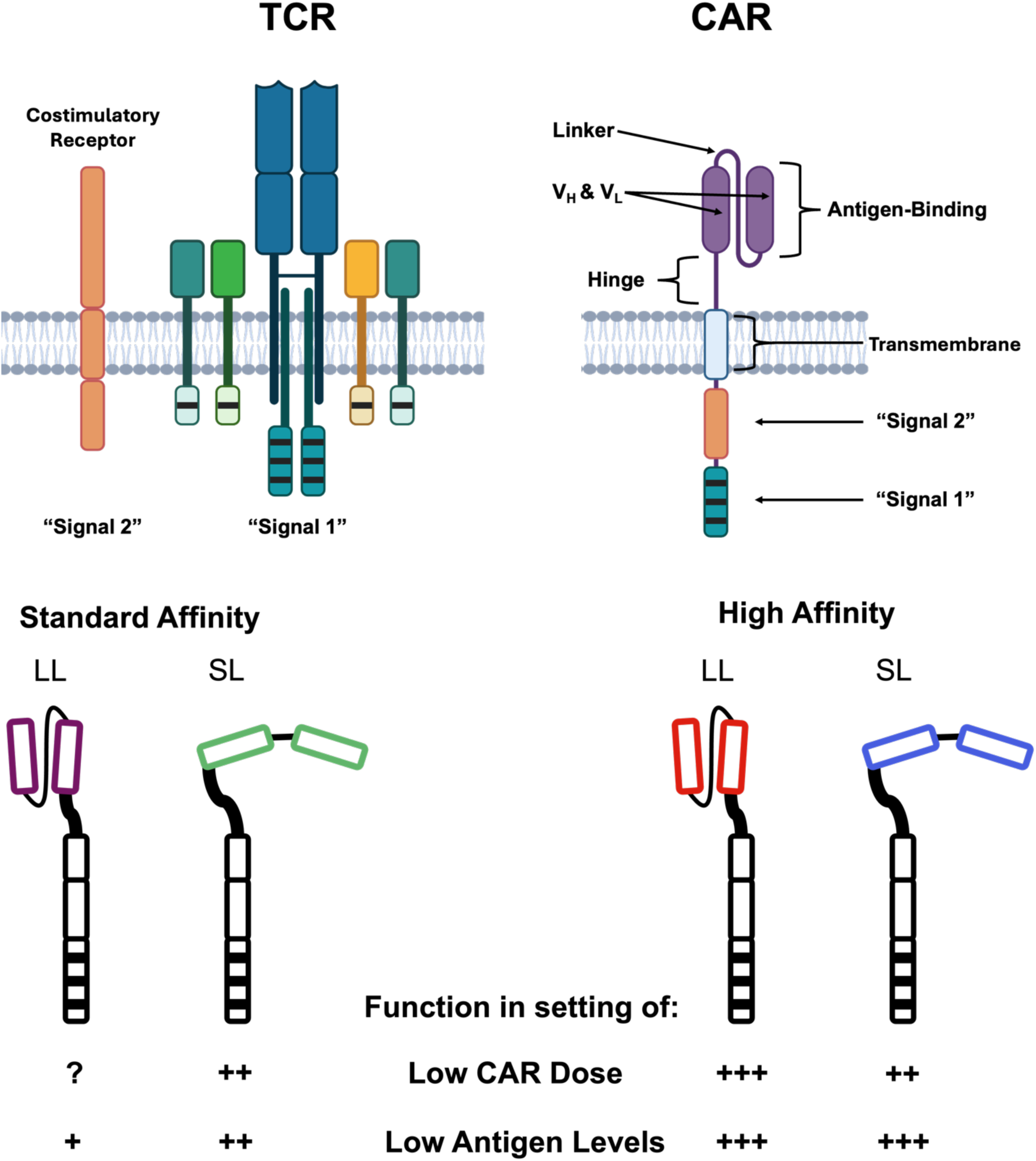

## INTRODUCTION

B-lineage acute lymphoblastic leukemia (B-ALL) is the most common type of pediatric malignancy, comprising nearly one-third of all pediatric cancers^1,2^. While T cells bearing chimeric antigen receptors targeting CD19 (CD19-CAR) have been highly successful in treating relapsed and/or treatment-refractory (r/r) disease. Remarkably, CD19-CAR therapy induces remissions in greater than 90% of patients in some trials^3^. Despite extraordinary initial successes, approximately 50% of patients treated with CAR T cells relapse after therapy, commonly due to CD19 downregulation or alternative splicing^4^. Therefore, targeting additional B-lineage antigens in multi-targeting modalities or follow up CAR T cell infusions has the potential to further improve patient outcomes. While multiple CD19-CAR products have now been FDA-approved for B-ALL, approval for CARs targeting additional antigens have been limited to B cell maturation antigen (BCMA), another B-lineage antigen with more consistent expression on multiple myeloma^3^. The CD22 antigen represents a highly favorable target antigen to build beyond the successes seen with CD19-CAR. Indeed, CD22-targeted CAR T cells (CD22-CAR) have demonstrated promising results in clinical trials for patients with relapsed and/or treatment-refractory B-ALL, including those resistant to CD19-targeted CAR therapy, but maintaining durable remissions remains a challenge. Recent follow-up data from the ongoing CD22-CAR trial at the NCI indicates that among patients who initially achieved remission, relapse-free survival is only 25% at 10 months post-CAR, and CD22^Lo^ relapse is common. Furthermore, patients who achieved MRD-negative CR had a significantly higher initial leukemic CD22 expression than patients who did not^5, 6, 7^. Together, these findings suggested that downmodulation of CD22 expression is a mechanism of leukemic resistance to CD22-CAR therapy in the clinic. This was corroborated by multiple pre-clinical studies with CARs targeting different hematologic target antigens, including CD19, CD20 and CD22 as well as solid tumor antigens like ALK and GPC2, which have shown that CAR T cells are less effective in the setting of decreased target antigen density^6, 8, 9, 10, 11^. These findings highlight the importance of developing strategies to improve both the sensitivity and durability of CAR efficacy in response to challenging targets such as CD22, which has a broad range of expression levels across patients and may be downmodulated in response to CAR pressure^5, 6, 7^. While direct modulation of target antigen density has proven to be a feasible strategy to boost efficacy of the CD22-CAR in preclinical models^6^, modifying structural components of the CAR such as antigen binding domain may be a more universally feasible strategy to boost CAR responses.

A component of CAR T cell biology highly linked to antigen binding domain is the intensity of constitutive antigen-independent signaling in the T cell, termed “tonic signaling”. Prior work demonstrated that an affinity-matured anti-GD2 binding domain drove strong tonic signaling, producing a functional, phenotypic and transcriptomic profile consistent with T cell exhaustion, which was improved by incorporating a 4-1BB costimulatory domain rather than CD28^12^. In contrast, tonic signaling was found to be essential for clinical responses in the context of the CD22-CAR. The NCI trial was the first to test the CD22-CAR and saw a complete remission rate exceeding 70%^5^. Trials at other institutions used nearly identical CARs with the same antigen binding domain from the m971 antibody clone, but swapped the flexible Glycine-Serine linker between antibody heavy and light chain from the “short” [G_4_S]_x1_ linker used at the NCI to a more standard “long” linker consisting of 3 to 4 tandem [G_4_S] motifs. Despite the minimal difference in CAR construct, survival of patients in these trials was drastically different, with the long linker CAR inducing complete remissions in only about 50% of patients, depending on the trial ^5, 13, 14^ ^51, 52^. Comparative laboratory studies found that the short linker significantly alters CAR biology, facilitating antigen-independent clustering of the chimeric receptor on the T cell surface, increasing tonic signaling, and increasing T cell activation as compared to the long linker CAR, seemingly augmenting T cell functionality. However, in other studies, the short linker CD22-CAR showed a similar phenotypic profile to the highly dysfunctional anti-GD2 CAR during *in vitro* expansion, indicating potential for T cell dysfunction in the long term (albeit with partial “rescue” via 4-1BB costimulation)^15,14^. Indeed, it has now become accepted at multiple institutions that the potential for dysfunction mediated by a short linker is necessary for maximizing the clinical efficacy of the CD22-CAR. While adjustment of binding affinity has also been explored as a method to enhance CAR efficacy in other CARs, functional effect has been highly dependent on CAR construct and target antigen biology^16,17,18, 19, 20 21.^

Given the importance of improving CAR responses to CD22 in B-ALL and ample evidence linking antigen binding domain design to CAR function and antitumor efficacy, we sought to rationally modify the antigen binding domain to understand whether we could maintain antigen sensitivity while removing the requirement for tonic signaling. To uncouple the impacts of CAR affinity and tonic signaling, we generated CARs with standard affinity (SA) or an m971-derived affinity matured (HA) binding domain, bearing either short linker (SL) or long linker (LL). We reveal that the standard affinity/long linker CAR (termed SA-LL), shown to be less clinically effective, has equivalent *in vitro* and *in vivo* function against WT leukemia to the standard affinity/short linker CAR (SA-SL), but specifically lacks function against CD22^Lo^ leukemia. Based on these findings, we hypothesized that for CD22-CAR, the binding affinity could be increased to specifically boost response to antigen-low leukemia, and that a long linker could be incorporated to reduce tonic-signaling-mediated dysfunction. We found that CARs incorporating the HA binder with either SL or LL format showed consistent improvement in early clearance of CD22^Lo^ leukemia and extension of survival *in vivo*. However, at a limiting CAR+ cell dose (mimicking high tumor burden), the HA-LL CAR improved clearance of WT leukemia relative to the SA-SL or HA-SL CAR, while extending survival at a higher CAR dose, indicating that a LL may allow for resistance to dysfunction relative to tonically-signaling SL CARs. These findings underscore the influence of multiple properties of antigen binding domain on anti-leukemic efficacy in the setting of clinically relevant relapse modalities and indicate that an increased affinity binder abrogates the requirement for tonic signaling in the CD22 CAR, allowing for resistance to dysfunction at a low CAR dose, while maintaining the ability to respond to antigen-low leukemia.

## RESULTS

### Antigen density impacts *in vitro* cytotoxicity and activation and *in vivo* leukemia clearance of CD22-CAR T cells

Prior studies by our lab and others have shown that target antigen density can have a profound impact on CARs targeting a variety of antigens, reducing *in vitro* effector functions and *in vivo* anti-tumor efficacy^6, 8, 9, 10, 11^. We set out to explore the impacts of antigen density on T cells expressing the original clinically-validated CD22-CAR (SA-SL) using NALM6 clones engineered to express varying levels of CD22^5^. *In vivo*, the standard CAR almost entirely cleared parental Nalm6 leukemia by day 8; however, while the CAR did show some efficacy in delaying the progression of CD22^Lo^ leukemia relative to CD22^Neg^ leukemia, the CD22^Lo^ leukemia continued to progress (Figures 1A-B). To further interrogate the impacts of CD22 density on CAR T cell functions, we tested this CAR in multiple *in vitro* assays across CD22 antigen densities. CAR T cell degranulation was directly impacted by antigen density, both in assays measuring bulk T cell function (Granzyme B ELISA) and on a single cell basis (flow cytometry for CD107a expression) (Figures 1C-D). Additionally, classical cell-surface markers of T cell activation including CD69 and CD25 were upregulated to a lesser extent in CARs co-cultured with CD22^Lo^ leukemia as compared to WT or CD22^Hi^ variants (Figures 1E-F). Together, these data demonstrate that low target antigen density negatively impacts multiple facets of *in vitro* and *in vivo* anti-CD22 CAR T cell activation and anti-tumor functions.

**Figure 1:**
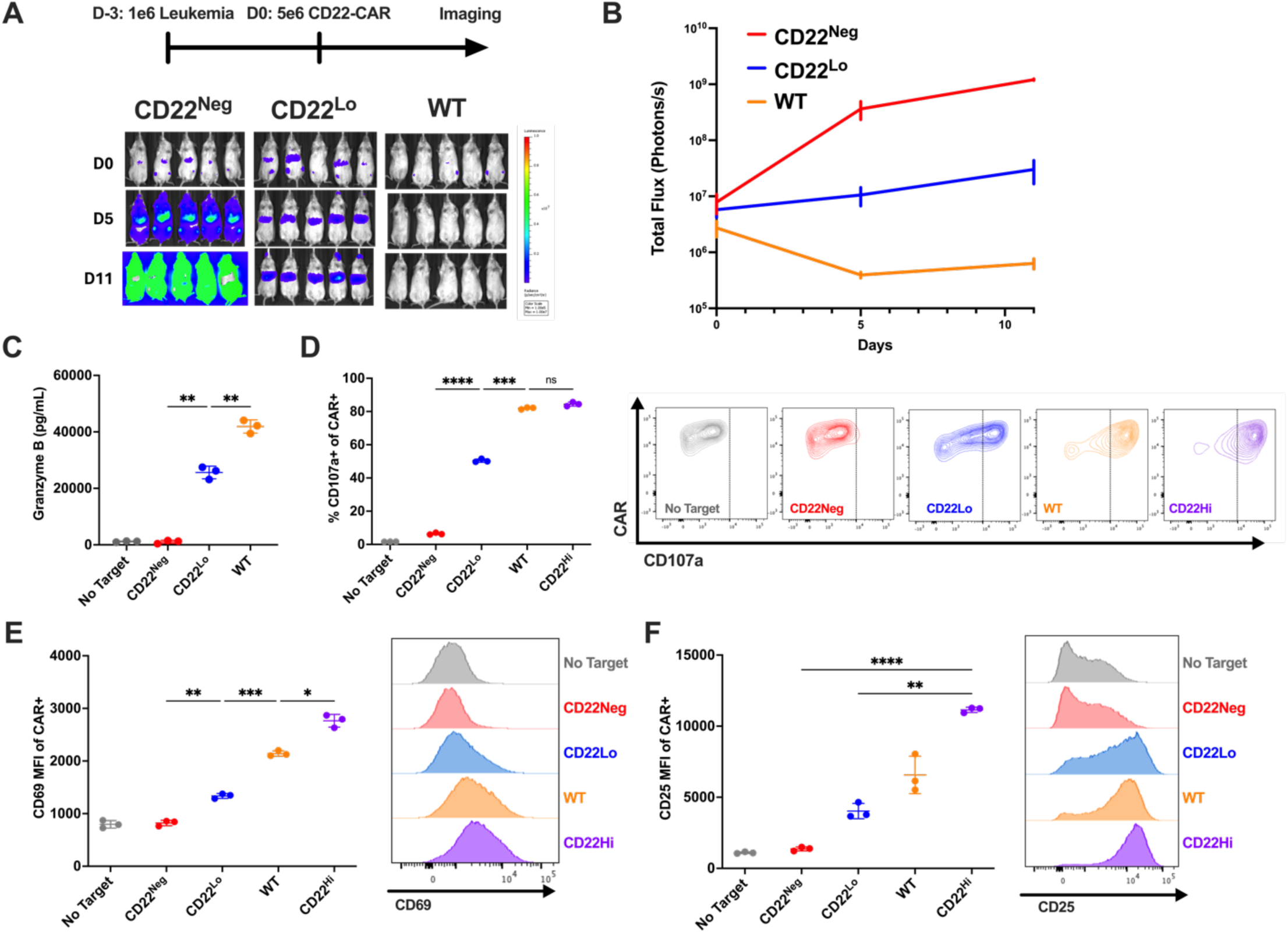
Antigen density impacts *in vitro* cytotoxicity and activation and *in vivo* leukemia clearance of CD22-CAR T cells. **1A:** Schematic: Timeline for *in vivo* experiment. NSG mice were injected with 1e6 indicated Nalm6 leukemia on day -3, followed by 5e6 CD22-CAR T cells on day 0. Bioluminescent imaging was performed before CAR dosing on day 0, as well on days 5 and 11 post-CAR. **1B:** Quantification of bioluminescence data in A. **1C:** ELISA measuring Granzyme B in supernatant after 16 hour co-culture of CD22-CAR T cells with the indicated leukemia. **1D:** Degranulation as measured by CD107a expression after 4 hour co-culture assay. **1E:** Activation as measured by CD69 expression after 6 hour co-culture assay. **1F:** Activation as measured by CD25 expression after 24 hour co-culture assay. All *in vitro* assays performed with n=3 technical replicates, 1 experiment. *In vivo* assay performed with n=5 mice per group, 1 experiment. Data represent mean +/-SD. * p<0.05, ** p<0.01, *** p<0.001, **** p<0.0001.

### Short linker boosts function of standard affinity CAR T cells in the setting of CD22^Lo^ leukemia

Recently, several clinical trials showed disparate results between CD22-targeted CAR T cells used to target B-cell malignancies, with one of the major distinctions being only in the length of the flexible linker between the heavy and light chain in the CAR antigen-binding domain^5, 13, 14^ ^51, 52^. Given the impact of target antigen density on anti-CD22 CAR T cell function, and the reported mechanism of CD22-downregulation after CAR treatment, we wanted to understand the impact of linker length on CAR T cell function in the context of both WT and CD22^Lo^ leukemia, with the prediction that a short linker would enhance CAR T cell function especially in the setting of low target antigen density. To test our hypothesis, we generated a CAR differing only in the length of the flexible linker between the heavy and light chains of the m971 scFv, using a longer (G_4_S)_x3_ linker rather than the (G_4_S)_x1_ linker found in the original CAR. We adopt the naming scheme “standard affinity/short linker” (SA-SL) for the original CAR, and “standard affinity/long linker” (SA-LL) for the long linker variant. *In vitro*, we found that while the linker length had minimal impact on the proportions of cells producing IFNg or IL-2 in response to WT Nalm6 leukemia, the SA-SL CAR showed enhanced response against CD22^Lo^ Nalm6 (Figures 2A-B). While linker length showed no significant impact on *in vivo* tumor clearance or long-term survival of mice bearing parental Nalm6, SA-SL CAR T cells mediated enhanced leukemia clearance and survival of mice bearing Nalm6-CD22^Lo^ (Figures 2C-E). Therefore, we demonstrate that despite similar performance against leukemia with high antigen density, the short linker is necessary for enhanced *in vitro* and *in vivo* function against CD22^Lo^ leukemia, with proportions of CAR T cells producing cytokine *in vitro* correlating with *in vivo* efficacy against CD22^Lo^ leukemia.

**Figure 2:**
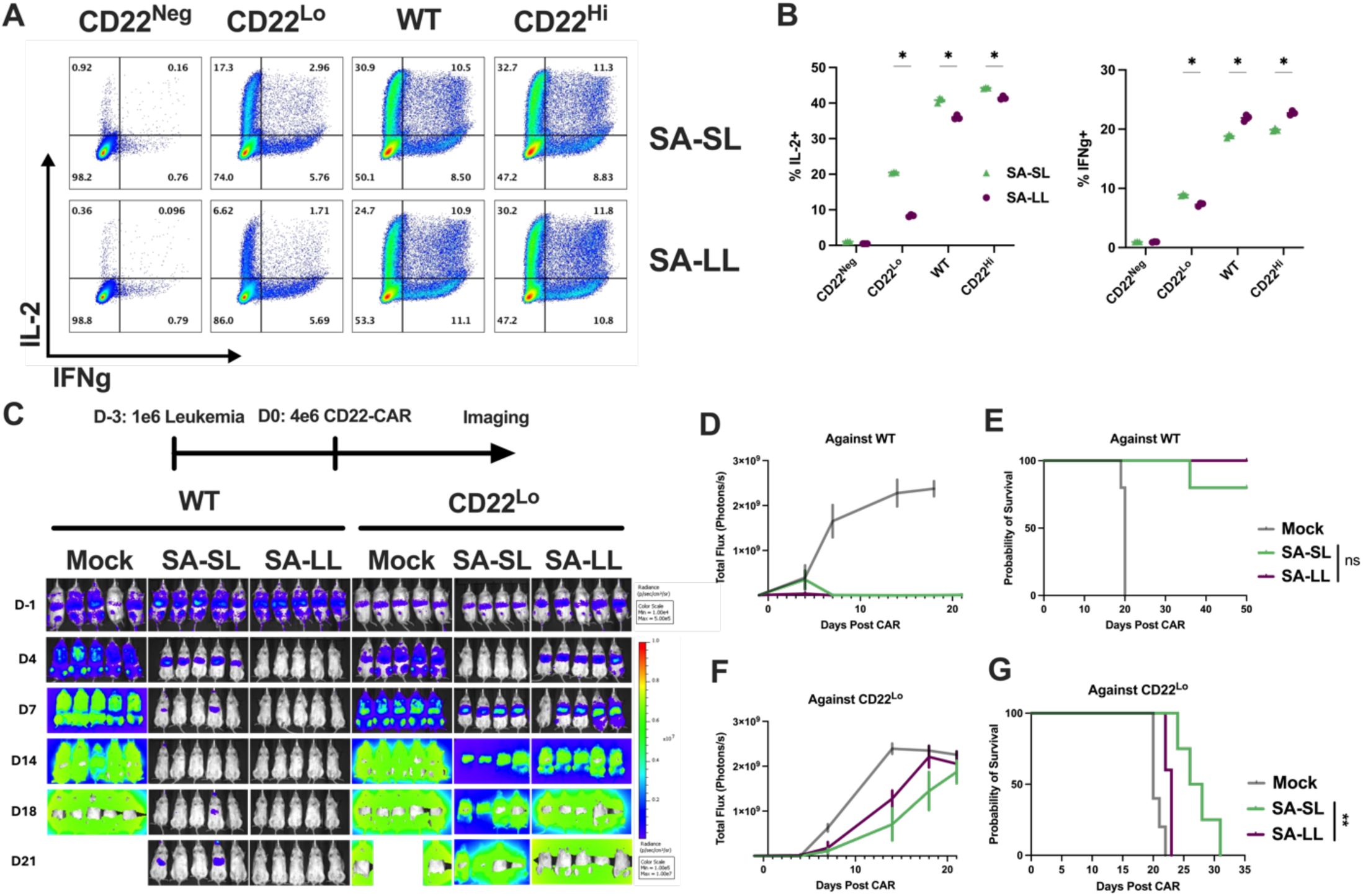
Short flexible linker boosts function of standard affinity CAR T cells in the setting of CD22^Lo^ but not WT leukemia. **2A:** Flow cytometry plots showing IL-2 by IFNg production after 6 hour coculture of the indicated CD22-CAR T cell with the indicated leukemia. **2B:** Quantification of cytokine data in A. **2C:** Schematic: Timeline for *in vivo* experiment. NSG mice were injected with 1e6 indicated Nalm6 leukemia on day -3, followed by 4e6 CD22-CAR T cells on day 0. Bioluminescent imaging was performed before CAR dosing on day -1, and biweekly post-CAR injection. Mice were monitored for survival. **2D:** Quantification of bioluminescence data against WT leukemia from C. **2E:** Survival of mice bearing WT leukemia. **2D:** Quantification of bioluminescence data against CD22^Lo^ leukemia from C. **2E:** Survival of mice bearing CD22^Lo^ leukemia. All *in vitro* assays performed with n=3 technical replicates, and are representative of two experiments with two independent donors. *In vivo* assay performed with n=5 mice per group, 1 experiment. Data represent mean +/-SD. * p<0.05, ** p<0.01, *** p<0.001, **** p<0.0001.

### A long linker enhances cytokine production by the high affinity CD22-CAR and abrogates persistent expansion

Functional disparities in the short and long linker m971-based CARs have been attributed to antigen-independent clustering which drives tonic signaling of the SA-SL CAR at baseline, characteristics not present in the SA-LL CAR ^14^. In Figure 2, we showed that in the context of the CD22 CAR, tonic signaling enhances anti-tumor responses against CD22^Lo^ leukemia. However, the tonically signaling GD2-CAR drives T cell dysfunction in the long term and has a basal transcriptional program consistent with T cell exhaustion. These characteristics were accompanied by a basal phenotypic state characterized by expression of multiple coinhibitory receptors, a profile shared by the CD22-CAR^15^. We hypothesized that a higher affinity antigen binding domain could maintain anti-tumor efficacy against CD22^Lo^ leukemia while mitigating the need for tonic signaling driven by a short linker, allowing for enhanced potency and resistance to dysfunction in the setting of parental leukemia. We previously used error prone PCR mutagenesis and yeast display followed by serial bio-panning over immobilized CD22 antigen to produce a m971-derived scFv with approximately 1000-fold higher affinity, and validated a CAR construct with this binder^6^. However, the high affinity CAR previously tested contained a more standardized long (G_4_S)_x3_ linker (“HA-LL”) rather than the short (G_4_S)_x1_ linker found in the original m971 CAR, a disparity potentially confounding the comparison of these two constructs. To explore the interplay between affinity and linker length, we generated the final combination of a high affinity binder with a short linker (“HA-SL”).

As the SA-LL CAR was shown to be inferior to the SA-SL CAR *in vitro*, *in vivo* and in multiple clinical trials, we chose to benchmark the HA CARs against the SA-SL CAR for all further *in vitro* and *in vivo* testing. To validate the enhanced affinity of the HA CARs relative to the SA, we transduced SA-SL, HA-SL and HA-LL CARs into T cells and performed a direct flow-based antigen binding assay with fluorophore-conjugated CD22 protein Fc. Indeed, both high affinity CARs showed increased affinity relative to the standard affinity CAR, as indicated by increased MFI at low antigen concentrations (Figure S1B).

To validate *in vitro* functionality of the affinity-matured CARs, we utilized the same *in vitro* intracellular cytokine staining (ICCS) assay shown to associate with *in vivo* CAR T cell efficacy in the standard affinity CAR T cells. All three CARs showed function against the CD22 target antigen. Against WT leukemia we observed that the SA-SL CAR had higher proportions of cells producing IL-2, IFNg, or both cytokines as compared to either high affinity CAR (Figures S2A-C). However, against CD22^Lo^ leukemia, the HA-LL CAR had significantly higher proportions of CAR+ cells producing cytokines as compared to HA-SL or SA-SL CARs (Figures S2D-F). Interestingly, the HA-LL CAR also had much greater preservation of proportions of cells producing cytokines with the drop in antigen density, as compared to the other two CARs. All CARs had very low background cytokine production against CD22^Neg^ leukemia, confirming the antigen-specific activity of these receptors (Figures S2G-H). To compare *in vitro* expansion potential of the CARs, we used a coculture assay with an internal fluorescent counting bead control in each well to quantify precise ratios of CAR:Beads and Leukemia:Beads. All three CARs expanded and cleared leukemia equivalently against WT Nalm6 (Figure S2I-J). Interestingly, while all three CARs also cleared CD22^Lo^ leukemia equivalently, both CARs with short linkers continued to expand after leukemia clearance, while the HA-LL CAR contracted post-antigen clearance, similar to a standard T cell response (Figure S2K-L). As the HA-LL CAR had been previously validated^6^, these data established that the HA-SL CAR was functional in the high affinity format and provided rationale for comparing all three CARs *in vivo*. Additionally, we demonstrate that in contrast to the standard affinity CARs compared in Figure 2, a long linker mediates superior *in vitro* cytokine production against CD22^Lo^ leukemia for the high affinity binder.

### HA-LL CAR mediates enhanced *in vivo* leukemia clearance against WT leukemia at limiting CAR+ cell dose and extends survival at high dose

We next wanted to test whether a short linker drove dysfunction in the setting of a limiting CAR+ cell dose and high target antigen expression. To test this, we again compared the high-affinity CAR constructs (HA-SL and HA-LL) against the SA-SL CAR, using a low 2e6 CAR dose to treat WT Nalm6 leukemia *in vivo*. At this dose, the high affinity CARs both outperformed the SA-SL CAR in early clearance of leukemia, with the HA-LL CAR showing very rapid and robust clearance of leukemia by day 4, while the other CARs showed highly variable leukemia burdens through day 11. While all CARs cleared leukemia by day 11, we found that the HA-LL CAR showed the most durable clearance of leukemia, while groups treated with the SA-SL and HA-SL CARs began to relapse at day 18 (Figures 3A-C). These results are consistent with our previous findings that the HA-LL CAR skewed T cells toward less-differentiated phenotypes based on cell-surface markers, which may allow for resistance to dysfunction^6^. Interestingly, while early *in vivo* tumor burden of WT Nalm6 with a higher but still subcurative 4e6 CAR+ cell dose was highly variable (Figures 3D-E), long-term survival was significantly extended in mice treated with HA-LL CAR T cells (Figure 3F). We demonstrate here that under stress conditions of sub-curative CAR+ cell doses, the HA-LL CAR shows superior clearance and extended survival in mice bearing high-antigen expressing WT leukemia, consistent with an enhanced ability to mediate leukemia clearance in the setting of high tumor burden.

**Figure 3:**
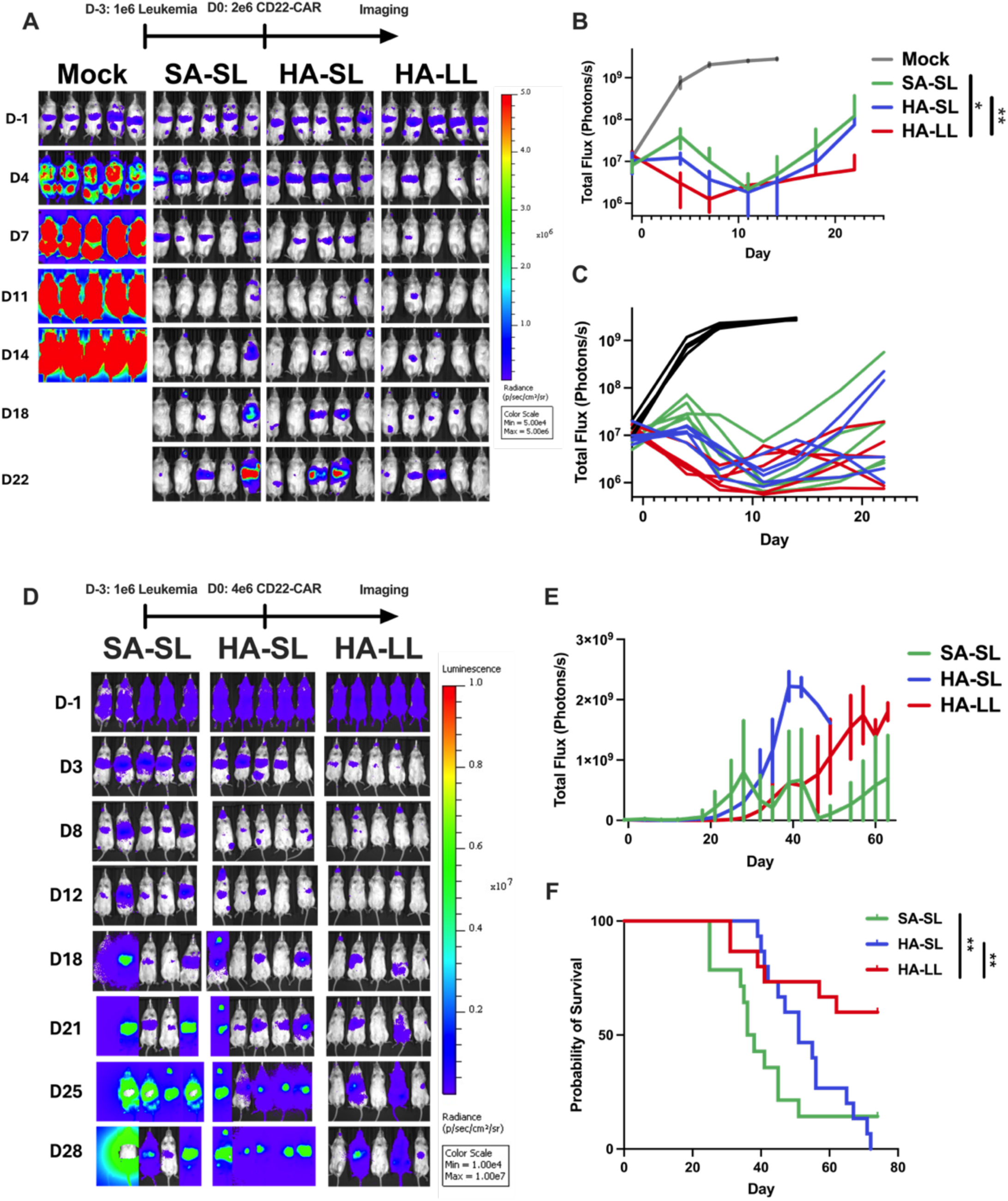
HA-LL CAR mediates enhanced *in vivo* leukemia clearance against WT leukemia at limiting CAR+ cell dose and extends survival at high dose. **3A:** Schematic: Timeline for *in vivo* experiment. NSG mice were injected with 1e6 WT Nalm6 leukemia on day-3, followed by 2e6 of indicated CD22-CAR T cells on day 0. Bioluminescent imaging was performed before CAR dosing on day -1, and biweekly post-CAR. **3B:** Quantification of average bioluminescence data for each group in A. **3C:** Quantification of individual bioluminescence data for each group in A. **3D:** Schematic: Timeline for *in vivo* experiment. NSG mice were injected with 1e6 WT Nalm6 leukemia on day -3, followed by 4e6 of indicated CD22-CAR T cells on day 0. Bioluminescent imaging was performed before CAR dosing on day -1, and biweekly post-CAR. **3E:** Quantification of average bioluminescence data for each group in D. **3F:** Survival of mice treated with 4e6 of the indicated CAR T cells. *In vivo* assay performed with n=5 mice per group, 1 experiment (3A to 3C) or 3 experiments with independent donors (3D to 3F). 3D to 3E are representative data from one experiment. 3F is pooled data, SA-SL (n=15), HA-SL (n=15), HA-LL (n=15). Data represent mean +/- SD. * p<0.05, ** p<0.01, *** p<0.001, **** p<0.0001.

### HA-LL CAR confers enhanced *in vivo* leukemia clearance and extends survival against CD22^Lo^ leukemia

Based on our results in Figure 1, SA-LL CAR T cells showed reduced function relative to SA-SL in the setting of CD22^Lo^ leukemia, a known clinical relapse mechanism in the setting of the CD22 target antigen, which could contribute to the poor clinical outcomes seen with long linker CARs ^5^. To test our hypothesis that boosting CAR affinity could eliminate the requirement for a short linker in driving *in vivo* CAR T cell function in response to leukemia with low levels of target antigen, we tested short and long linker high affinity CARs (HA-SL and HA-LL) against the SA-SL CAR in our xenograft model against CD22^Lo^ leukemia (Figure 4A). Across four T cell donors, we found that both high affinity CARs consistently outperformed the SA-SL CAR, mediating enhanced tumor clearance by bioluminescence imaging and flow cytometry of bone marrow at day 17-19 post CAR treatment (Figure 4A-C). Interestingly, in contrast to the standard affinity CAR, this was not dependent on the linker length, and both high affinity CARs performed equivalently. Despite consistently higher leukemia burden, the SA-SL CAR showed the highest proportions of CAR+ cells in the marrow, indicating that the remaining CAR T cell population was likely in a dysfunctional state (Figure 4D-E). While both high affinity CARs performed similarly early on, only the long linker CAR significantly extended survival against CD22^Lo^ leukemia (Figure 4F). As a whole, these data demonstrate that affinity maturation of antigen binding domain, combined with a long linker, enhances CAR T cell function against CD22^Lo^ leukemia. These results are in contrast to standard affinity CARs, which necessitate a short linker for function against CD22^Lo^ leukemia.

**Figure 4:**
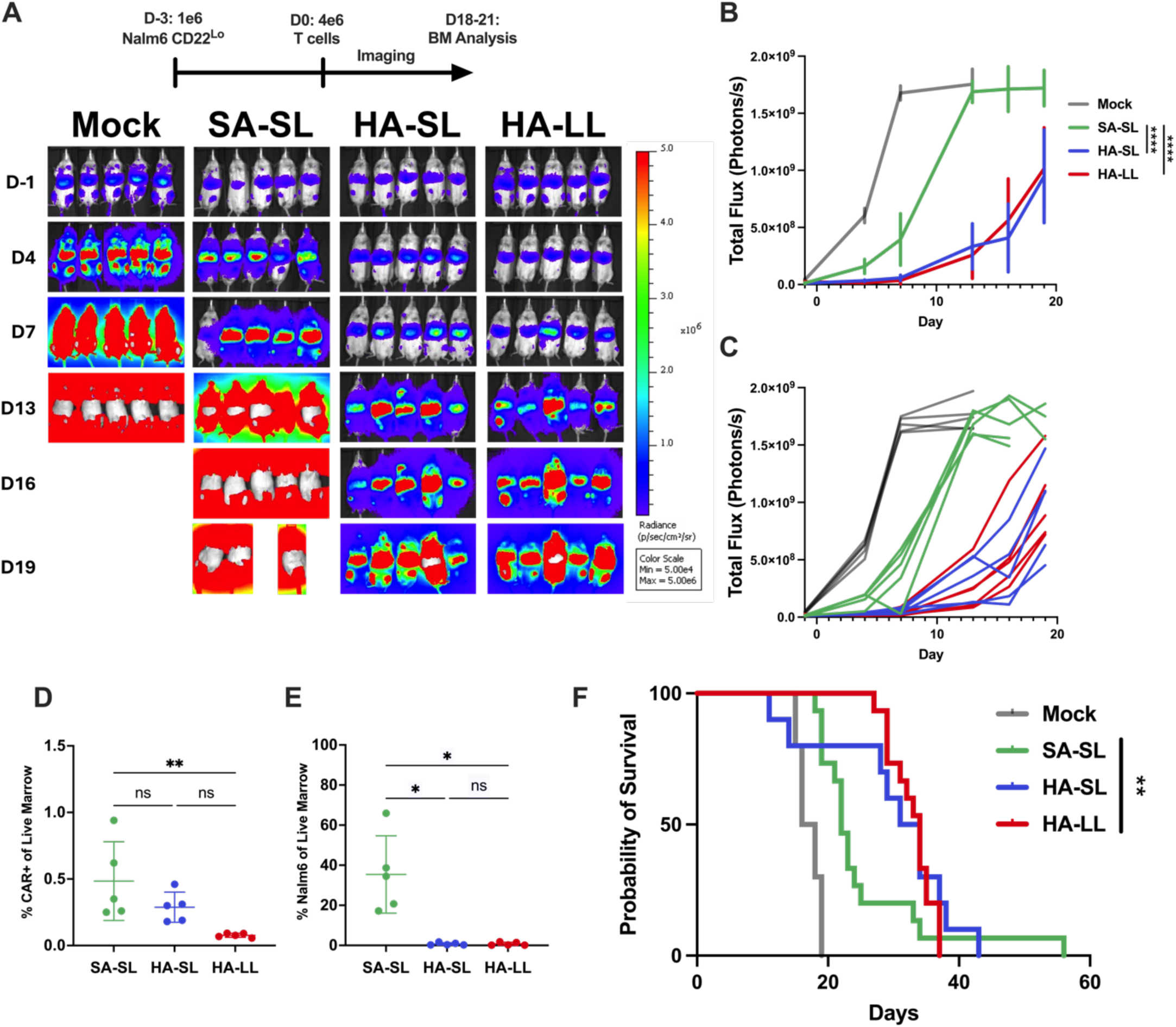
HA-LL CAR confers enhanced *in vivo* leukemia clearance and extends survival against CD22^Lo^ leukemia. **4A:** Schematic: Timeline for *in vivo* experiment. NSG mice were injected with 1e6 CD22^Lo^ Nalm6 leukemia on day -3, followed by 4e6 of indicated CD22-CAR T cells on day 0. Bioluminescent imaging was performed before CAR dosing on day -1, and biweekly post-CAR. **4B:** Quantification of average bioluminescence data for each group in A. **5C:** Quantification of individual bioluminescence data for each group in A. For 4D to 4E, bone marrow was analyzed by flow cytometry at day 18 post-CAR for indicated cell population. **4D:** % CAR+ of live marrow. **4E:** % leukemia of live marrow. **4F:** Survival of mice treated with 4e6 of indicated CAR T cells. *In vivo* assay performed with n=5 mice per group, 4 experiments with independent donors. Data in 4A to 4C is representative data from one experiment. Survival is pooled from 3 experiments with independent donors: Mock (n=10), SA-SL (n=15), HA-SL (n=10), HA-LL (n=15). Data represent mean +/- SD. * p<0.05, ** p<0.01, *** p<0.001, ****

## DISCUSSION

CAR T cells targeted to the B-lineage antigen CD19 have revolutionized the therapy of B-lineage malignancies such as acute lymphoblastic leukemia, resulting in remission rates of 70-90% when used to treat relapsed and treatment-refractory disease. However, 50% of patients relapse, often with CD19^Neg^ disease no longer targetable by CD19-CAR T cells^4^. Therefore, other strategies to confer more durable remissions in higher proportions of patients are critical. One successful intervention in treating CD19^Neg^ disease is use of CAR T cells targeted to another B-lineage antigen, CD22. While CD22-CAR T cells have a 70% rate of remission induction, even when treating CD19^Neg^ disease, patients still relapse and commonly present with leukemia displaying low surface expression of the CD22 antigen relative to pre-treatment levels ^5^. Therefore, while creating a CAR with high sensitivity to target antigen is essential for extending durability of remissions, the already clinically effective CD22-CAR has been shown to rely on antigen-independent CAR clustering and tonic signaling for efficacy, a feature which has been shown to negatively impact T cell function in other settings^15, 22^. This tonic signaling, driven by a short (G_4_S)_x1_ linker in the scFv, has been essential for strong clinical responses in patients, despite similar efficacy of m971-based CARs with longer linkers in preclinical models^5, 13, 14^ ^51, 52^. Known tumor relapse modalities post-CAR therapy present a useful opportunity for understanding the biology of these artificial antigen receptors and the T cells bearing them. Here, we have presented data from experimental systems designed to model the known relapse modalities of antigen downregulation/heterogeneity (leukemia engineered to express lower target antigen density) and “exhaustion-like” T cell dysfunction (administration of a subcurative CAR+ cell dose). Here, we show that while standard affinity CAR T cells with long (G_4_S)_x3_ linkers can mediate strong responses against high antigen-expressing WT leukemia in xenograft models, a short linker is required for the *in vivo* response against CD22^Lo^ leukemia. We build on this finding using a previously generated CD22-CAR construct with an antigen binding domain of approximately 1000-fold higher affinity, but here test short and long linker versions of this construct and compare these to short and long linker versions of the standard affinity CAR^6^. We find that a long linker allows for more durable *in vivo* functionality of the HA CAR in settings of limiting CAR dose. Finally, we show that the HA CAR does not require a short linker for activity against CD22^Lo^ leukemia, and enhances leukemic clearance across multiple donors relative to the original m971-SL CAR. Therefore, we have employed rational modification of the antigen binding domain in CD22-targeted CAR constructs to design a CAR which maintains the high antigen sensitivity required to successfully target the CD22 antigen, while alleviating the requirement for tonic signaling and the associated T cell dysfunction driven by the tonic signaling SA-SL CAR.

Many questions remain regarding characteristics of antigen binding domain and how they impact CAR T cell biology. While phosphoproteomic studies by Singh et al. indicated that linker length also dictated tonic signaling profile of a CD33-targeted CAR, the same was not true for a CAR targeting CD19, accompanying functional studies were not performed, and binding affinity was not considered for these antigens^14^. Therefore, one important avenue is broadly characterizing the interplay of affinity and linker length in CARs targeting other antigens, or with other receptor modifications such as alternative costimulatory domains. While tonic signaling would seem to lack utility in a solid tumor microenvironment characterized by high antigen burden and repetitive stimulation, a tonically signaling GD2-targeted CAR has been tested against solid tumors in multiple studies and comparisons of CARs targeting other solid tumor antigens are warranted ^15, 22, 23^. While our studies provide novel insight into the biology of the chimeric receptor in a bulk cell population, we and others have previously shown key differences in the biology of memory versus naïve-derived CAR T cells^24^, and between CD4 and CD8 CAR T cells^25^, suggesting a differential role for antigen receptor signal strength even within these canonical T cell subgroups. As an exploratory study, deep phenotypic comparisons using spectral flow cytometry or mass cytometry, or single cell RNA sequencing, could be performed to better characterize what features of T cell state are dictated primarily by linker length versus primarily by binding affinity. Other potential modifications of the antigen binding domain, such as using a heavy-chain only binding domain or targeting a different epitope on the relatively large CD22 molecule were not investigated here and should be an emphasis of future work. Finally, recent findings have shown that the CD22-CAR can cause a severe hyperinflammatory syndrome toxicity similar to classically defined hemophagocytic lymphohistiocytosis (HLH), which has been found to be highly prevalent in patients treated with the original tonically-signaling SA-SL CAR, and is distinct in presentation from cytokine release syndrome seen with other CAR T cells to date. This toxicity is in contrast to those commonly observed with other non-tonically signaling CARs such as the CD19-CAR, and may be mitigated by using a long-linker CAR more similar to the CD19-CAR, such as the HA-LL CAR described here. It is difficult to test exact mechanisms of HLH manifestations in pre-clinical models with human CAR T cells due to the complex immune interactions leading to this pathology, which are not present in immunodeficient NSG hosts. However, we observe that the HA-LL CAR rectifies increased expansion and delayed contraction, which were implicated in a recent clinical correlative study as drivers of HLH in CD22-CAR patients treated with the same SA-SL CAR that showed these characteristics in our studies^26^. An investigation into whether a tonic signaling CAR drives HLH-like toxicity in an immune-intact host would likely provide useful insight into the pathogenesis.

In conclusion, our work here has tested a non-tonic signaling affinity-matured CD22 CAR based around the same clinically-validated m971 antibody clone used in the standard affinity CAR. This CAR has the potential to directly improve outcomes of patients with B-lineage malignancies by resisting the known relapse mechanisms of antigen downregulation and T cell dysfunction, and potentially circumventing toxicities associated with the current CD22 CAR. CARs are highly modular molecules and have been readily modified to target an extensive set of antigens. Our evaluation as to how varying antigen binding affinity and tonic signaling impact CAR T cell biology may be clinically applicable to CARs targeting many other tumor types. As CAR T cells are now being used to treat SLE, multiple sclerosis, aging/cellular senescence, and to target virally expressed surface proteins (HIV)^27^, we expect these findings to also prove widely applicable toward the design of future CAR variants to mitigate a broad array of disease pathologies.

## MATERIALS AND METHODS

### Culture of cell lines and preparation of human donor T cells

The parental Nalm6 (GFP/Luciferase-transduced) B-ALL cell line was obtained from Dr. Crystal Mackall, Pediatric Oncology Branch, NCI, NIH, Bethesda, MD, 2008. Nalm6 CD22 site density model cell lines (CD22^Neg^, CD22^Lo^, CD22^Hi^) were previously generated as described ^5^. Briefly, CD22 was knocked out using CRISPR/Cas9, and a CD22 transgene was subsequently reintroduced using lentiviral transduction. Following transduction with CD22, single-cell cloning by limiting dilution was used to generate Nalm6 clones which stably express varying levels of CD22. CD22 site density was (antigen sites/cell) on Nalm6 clones was previously quantified using CD22-PE, and CD22 MFI was converted to sites/cell using QuantiBrite PE Beads (BD Biosciences) as previously described ^5^. Nalm6 WT and all CD22 site density clones were cultured in complete RPMI-1640 (cRPMI), RPMI-1640 medium (Gibco) supplemented with 10% heat-inactivated FBS (Omega Biosciences) and 2mmol/L L-glutamine (Gibco), at 37C and 8% CO_2_. The viral producer Lenti-X cell line was obtained from Takara Bio. Lenti-X cells were cultured in DMEM medium (Gibco) supplemented with 10% heat-inactivated FBS, 2mmol/L L-glutamine (Gibco), and 10mmol/L HEPES (Gibco), and were passaged at least once prior to transfection with lentiviral plasmids. All cell lines were routinely tested for *Mycoplasma* by the University of Colorado Anschutz core facility.

### Generation of human CD22 CAR constructs

For ease of cloning novel CAR variants, modular monovalent second-generation CAR constructs (4-1BB costimulatory domain with CD8 transmembrane) were previously synthesized (GeneArt, Thermo Fisher) and cloned into a lentiviral plasmid backbone. These constructs each incorporated restriction sites (XhoI and SpeI) flanking the scFv region of the CAR to allow for rapid synthesis of new CAR constructs. In the present study, variants of the m971 scFv were synthesized with flanking XhoI and SpeI restriction sites (GeneArt, Thermo Fisher) and subcloned into the CAR backbone using standard restriction enzyme cloning with XhoI and SpeI enzymes (New England BioLabs). CD22 CAR sequences were verified by Sanger sequencing (Eton Biosciences).

### Generation of human CD22 CAR T cells

Healthy human donor whole blood obtained from the Children’s Hospital Colorado Blood Donor Center, under an institutional board-approved protocol. Peripheral blood mononuclear cells were isolated using Lymphocyte Separation Medium (Corning) according to manufacturer protocol. T cells were then purified using an EasySep Human T Cell Isolation Kit (StemCell) and cryopreserved in 90% heat-inactivated FBS and 10% DMSO. Lentiviral vectors encoding CD22 CAR constructs were generated by transient transfection of the Lenti-X viral producer cell line with the CD22 CAR construct, as well as lentiviral packaging and envelope plasmids (pMDLg/pRRE, pMD.2G, and pRSV-Rev, all obtained from Addgene) using Lipofectamine-3000 (Thermo Fisher) in OptiMEM (Gibco). 6 hours following transfection, medium was replaced with fresh media. At 24 hours and 52 hours following transfection, supernatant containing lentiviral vector was harvested and spun at 3000 RPM for 10 minutes to remove cell debris and frozen at -80C for later use. Healthy human T cells were thawed at 1e6/mL in human T cell expansion media (hTCEM), AIMV medium (Gibco), supplemented with 5% heat-inactivated FBS, 2mmol/L L-glutamine, 10mmol/L HEPES, and 40 IU/ml recombinant human IL-2 (R&D Systems), and activated for 48 hours with CD3/CD28 Human T Activator Dynabeads (Gibco) at a ratio of 1:1 cells to beads. On days 2 and 3, T cells were transduced with lentivirus in the presence of 10 ug/mL protamine sulfate (Sigma) and 40 IU/mL IL-2 (R&D Systems), spinning for 2 hours at 1000xG. On day 4, CD3/CD28 Human T activator microbeads were removed using a magnetic rack and T cells were resuspended at 0.5e6/mL in hTCEM. T cells were expanded for 5-6 more days, with new media and cytokines added every other day. Following expansion, transduction efficiency of the CD22-CAR was evaluated by flow cytometry staining with CD22-Protein Fc (R&D Systems) and CAR T cells were cryopreserved or used immediately for *in vitro* or *in vivo* assays.

### Nalm6 xenograft *in vivo* studies

All xenograft studies were performed using NSG mice (NOD-scid-gamma, NOD.Cg-*Prkdc^scid^Il2rgtm1Wjl*/SzJ; Strain #: 005557, Jackson Laboratories). NSG mice received 1e6 GFP/Luciferase-positive leukemia cells (22^Neg^, 22^Lo^, or WT) via intravenous tail vein injection on Day -3. Mice then received CD22 CAR T cells via intravenous tail vein injection on Day 0 at the indicated dose level. Leukemia burden was measured twice weekly using Xenogen *In Vivo* Imaging System (IVIS) 200 or Spectrum (Caliper Life Sciences) after intraperitoneal injection with D-Luciferin (Perkin Elmer). Total flux (photons/s) was measured for each mouse using Living Image 4.7 (Caliper Life Sciences). All animal studies were conducted in accordance with an animal protocol approved by the University of Colorado Anschutz Medical Campus Institutional Animal Care and Use Committee (IACUC, Protocol 751).

### Flow cytometry for cell-surface and intracellular markers

Flow cytometric analysis of cell-surface and intracellular proteins was performed on a BD LSR Fortessa X-20 Flow Cytometer (BD Biosciences). For all CD22 CAR constructs, the CAR was detected by primary staining with a recombinant CD22-Fc chimera protein (R&D Systems) followed by secondary staining with PE-Conjugated Goat anti-Human IgG Fc secondary antibody (Thermo Fisher). The following human monoclonal antibodies were used to detect cell-surface proteins: CD3-APC/Cy7, CD3-BV421, CD45-PerCP/Cy5.5, CD8-APC, CD45RA-APC, CD25-BV711, CD107a-APC, CD69-BV605 (all from BioLegend). The following human monoclonal antibodies were used to detect intracellular phosphoproteins: cells were excluded using eBioscience Fixable Viability Dye eFluor 506 (Thermo Fisher/Invitrogen). GFP+ leukemia was identified using the FITC channel.

### *In vitro* assays for CD107a, CD69, CD25, intracellular cytokine staining and ELISA

*In vitro* assays were performed using a 1:1 effector to target cell ratio with 1e5 CD22 CAR T cells and 1e5 target Nalm6 leukemia cells (22^Neg^, 22^Lo^, WT, 22^Hi^) in cRPMI in 96-well plates followed by analysis by flow cytometry or ELISA at the indicated timepoints. For ELISA, following a 16 hour coculture, the plate was centrifuged at 300xg for 6 minutes to pellet cells and then supernatant was collected for analysis. Prior to analysis of GzmB production, supernatant was diluted 1:100 in RPMI before running the GzmB ELISA kit (Thermo Fisher) was then used to measure protein concentrations in the culture supernatant according to the manufacturer’s instructions. Final cytokine concentrations were calculated based on initial dilution factor. CD107a degranulation assays were performed by incubation for 4 hours in the presence of 2uM monensin and 1uL of CD107a antibody. ICCS was performed by incubation for 6 hours, with 1uM monensin and 2.5uM Brefeldin A added at 1 hour in. CD69 was analyzed after a 6 hour incubation. CD25 was analyzed after a 24 hour incubation.

### *In vitro* leukemia clearance and CAR T expansion assay

5e5 CD22 CAR T cells were co-cultured with 5e5 Nalm6 leukemia cells (22^Neg^, 22^Lo^, WT, 22^Hi^) and 1e5 inert AccuCheck Counting Beads (Thermo Fisher; Cat. #PCB100) in 1mL RPMI per well in a 12-well plate. At each timepoint, sample was taken from each well for flow cytometric analysis of CAR T cell expansion and leukemia clearance, and fresh RPMI media was added to each well to keep volume consistent. To determine relative cell quantities per sample, FlowJo v10 software (BD Biosciences) was used to gate on counting beads based on distinct SSC and FITC profile, and total counting beads per sample was counted and relative quantities of each cell type were represented as ratio of CAR T cells/Beads or Leukemia cells/Beads.

### Cell based direct antigen-binding affinity titration assay

CARs were stained with indicated concentrations of fluorophore-conjugated CD22 Protein Fc from 2ng/uL to 2000 ng/uL for HA CARs or 2 ng/uL to 4000 ng/uL for SA CAR (until antigen binding plateaued). MFI of CAR+ Populations were measured and normalized to peak antigen binding concentration for each individual CAR.

### Statistics

ELISA, CD107a, CD69 and CD25 data were compared using ordinary one-way ANOVA with Tukey’s multiple comparisons tests. Relative cell quantities in *in vitro* leukemia clearance and CAR T cell expansion experiments were compared using ordinary two-way ANOVA with Tukey’s multiple comparisons tests. ICCS data were compared using multiple unpaired t-tests with Benjamini, Krieger and Yekutieli procedure to control for false discovery rate, or ordinary one-way ANOVA with Tukey’s multiple comparison tests as appropriate for group numbers. *In vivo* flow cytometric analysis of % CAR T cells and leukemia burden data were compared using Kruskal-Wallis tests. For *in vivo* imaging studies, leukemia bioluminescence data were compared using a standard two-way ANOVA with post-hoc Tukey’s multiple comparisons tests. For comparison of survival curves, the Log-rank (Mantel-Cox) test was used. All statistical tests were run in Prism 10 software (GraphPad).

## DATA AVAILABILITY STATEMENT

All data is available from the corresponding author upon request.

## ACKNOWLEDGEMENTS

We acknowledge all members of the Fry and Kohler labs for help with bone marrow harvests and useful discussion regarding the manuscript.

## AUTHOR CONTRIBUTIONS

Conceptualization: K.R.D., Z.H.W., D.D., M.E.K., T.J.F.

Data curation: K.R.D, Z.H.W., L.L., C.B., A.N.

Formal analysis: K.R.D., Z.H.W.

Funding acquisition: D.D., T.J.F.

Investigation: K.R.D, Z.H.W., L.L., C.B., A.N., D.D.

Methodology: K.R.D., Z.H.W., M.E.K., T.J.F.

Project administration: L.L., M.E.K., T.J.F.

Resources: K.R.D, Z.H.W., L.L., C.B., A.N., D.D.

Software: K.R.D., Z.H.W.

Supervision: K.R.D., M.E.K., T.J.F

Validation: K.R.D., Z.H.W., T.J.F.

Visualization: K.R.D., Z.H.W.

Writing – original draft: K.R.D., Z.H.W., T.J.F.

Writing – review & editing: All authors.

## SUPPLEMENTAL FIGURES

**Figure S1:**
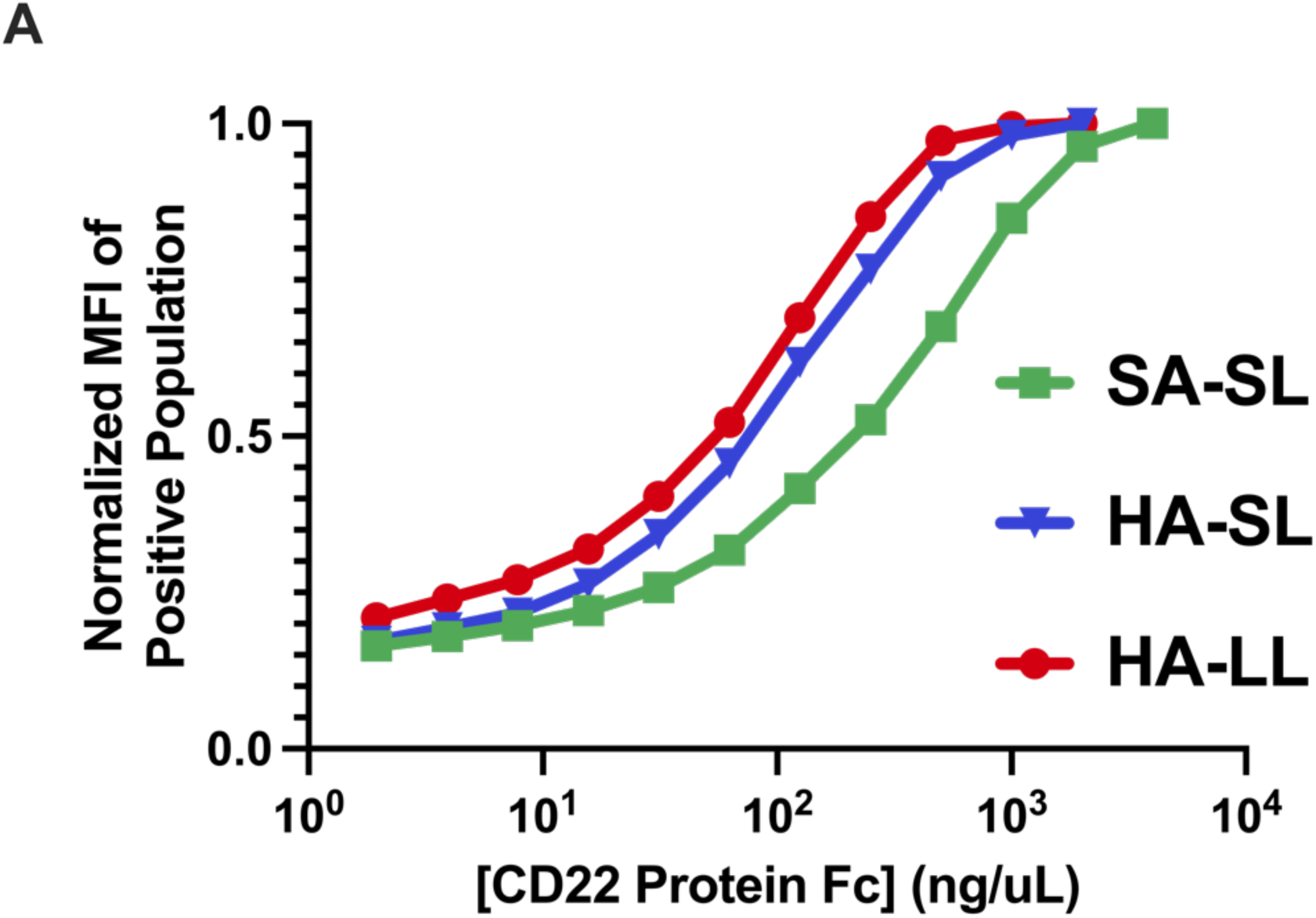
Affinity measurements of standard and high affinity binders in different formats. **S1A:** Cell-based direct antigen-binding affinity titration assay. Indicated CARs were stained with indicated concentrations of fluorophore-conjugated CD22 Protein Fc. MFI of CAR+ Populations were measured and normalized to peak protein binding for each individual CAR. Data represents one experiment with one replicate per concentration.

**Figure S2:**
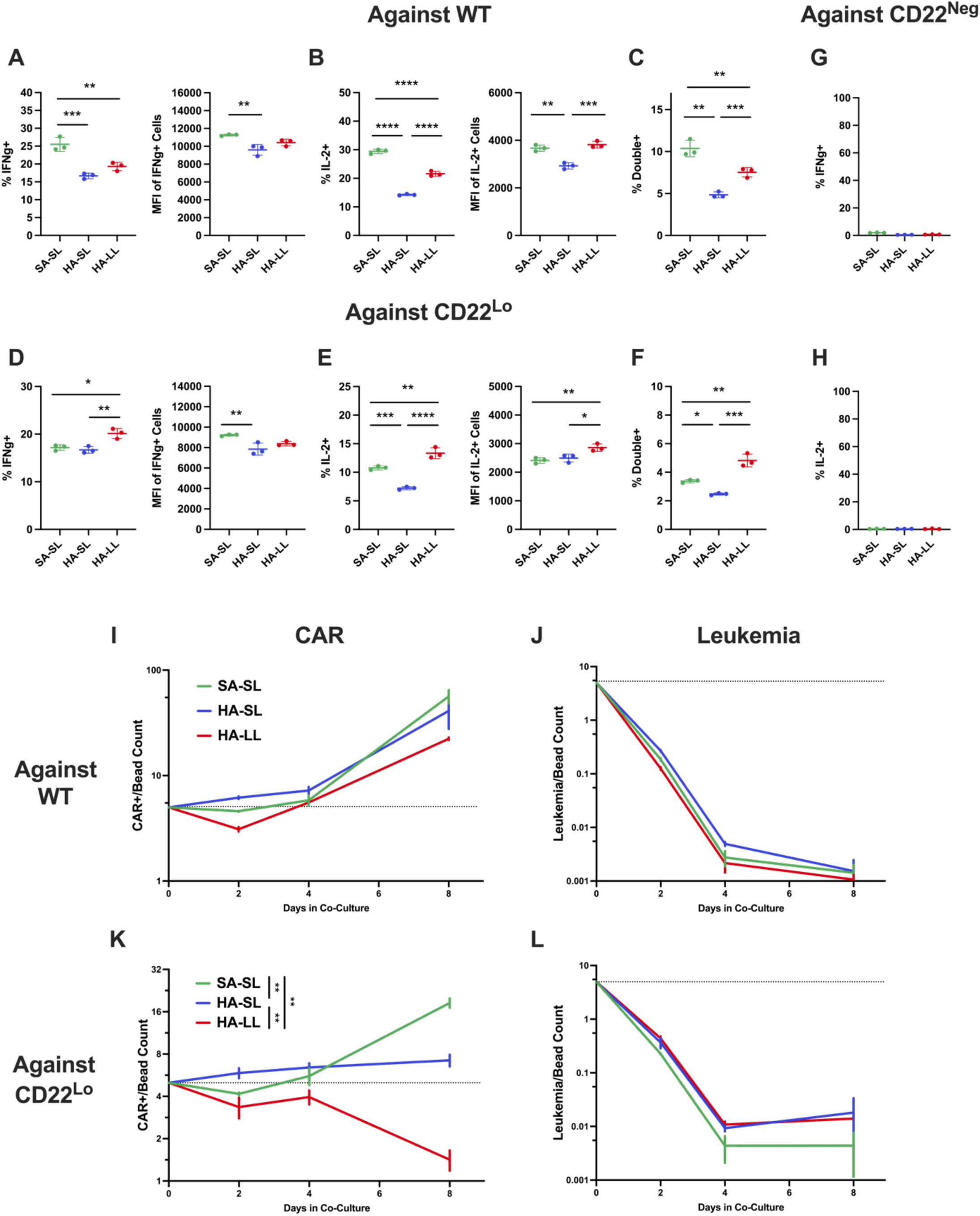
A long linker enhances cytokine production by the high affinity CD22-CAR. Figures S2A to S2F quantify indicated metrics by flow cytometry after coculture of indicated CD22-CAR with indicated leukemia after 6 hour coculture. **S2A:** %+ and MFI for IFNg production against WT leukemia. **S2B:** %+ and MFI for IL2 production against WT leukemia. **S2C:** %+ of cells making IFNg and IL-2 against WT leukemia. **S2D:** %+ and MFI for IFNg production against CD22^Lo^ leukemia. **S2E:** %+ and MFI for IL2 production against CD22^Lo^ leukemia. **S2F:** %+ of cells making IFNg and IL-2 against CD22^Lo^ leukemia. Figures S2G to S2J quantify CAR and leukemia counts relative to a starting 5:1 ratio of leukemia and CAR to fluorescent counting beads. Aliquots were taken from each condition and analyzed by flow cytometry at each of the indicated time points. **S2G:** Quantification of CAR Count/Bead Count ratio against WT leukemia. **S2H:** Quantification of Leukemia Count/Bead Count ratio for WT leukemia. **S2I:** Quantification of CAR Count/Bead Count ratio against CD22^Lo^ leukemia. **S2J:** Quantification of Leukemia Count/Bead Count ratio for CD22^Lo^ leukemia. All *in vitro* assays performed with n=3 technical replicates. Figure S2A-S2F are representative of two experiments with two independent donors. Figure S2G-S2J are representative of one experiment. Data represent mean +/- SD. * p<0.05, ** p<0.01, *** p<0.001, **** p<0.0001.

## TABLES

**Table 1.** Antibodies and Other Dyes

| Antibodies and Other<br>Dyes |  |  |  |  |  |
| --- | --- | --- | --- | --- | --- |
| Table 1 |  |  |  |  |  |
| Anti-Human | Specificity | Fluor | Company | Cat No | Clone |
|  | CD22 Protein Fc | N/A | R&D Systems | 1968-SL-050 | N/A |
|  | Human IgG Fc | PE | BioLegend | 409304 | HP6017 |
|  | CD107a | APC | BioLegend | 328620 | H4A3 |
| | IFNγ | BV711 | BioLegend | 502540 | 4S.B3 |
|  | IL-2 | BV421 | BioLegend | 500328 | MQ1-17H12 |
|  | CD3 | APC/Cy7 | BioLegend | 344818 | SK7 |
|  | CD3 | BV421 | BioLegend | 344834 | SK7 |
|  | CD45 | PerCP/Cy5.5 | BioLegend | 304028 | HI30 |
|  | CD25 | BV711 | BioLegend | 356138 | M-A251 |
|  | CD69 | BV605 | BioLegend | 310938 | FN50 |
| Other Dyes | Specificity | Fluor | Company | Cat No | Clone |
|  | Fixable Viability Dye | eFluor 780 | eBioscience | 65-0865-18 | N/A |
|  | Fixable Viability Dye | eFluor 506 | eBioscience | 65-0866-18 | N/A |
|  | CellTrace Violet | Violet 450 | Invitrogen | C34571 | N/A |

